# Thymic epithelial organoids mediate T cell development

**DOI:** 10.1101/2024.03.05.583513

**Authors:** Tania Hübscher, L. Francisco Lorenzo-Martín, Thomas Barthlott, Lucie Tillard, Jakob J. Langer, Paul Rouse, C. Clare Blackburn, Georg Holländer, Matthias P. Lutolf

## Abstract

**Although the advent of organoids opened unprecedented perspectives for basic and translational research, immune system-related organoids remain largely underdeveloped. Here we established organoids from the thymus, the lymphoid organ responsible for T cell development. We identified conditions enabling thymic epithelial progenitor cell proliferation and development into organoids with in vivo-like transcriptional profiles and diverse cell populations. Contrary to two-dimensional cultures, thymic epithelial organoids maintained thymus functionality in vitro and mediated physiological T cell development upon reaggregation with T cell progenitors. The reaggregates showed in vivo-like epithelial diversity and ability to attract T cell progenitors. Thymic epithelial organoids provide new opportunities to study TEC biology and T cell development in vitro, pave the way for future thymic regeneration strategies and are the first organoids originating from the stromal compartment of a lymphoid organ.**

**Summary statement:** **Establishment of organoids from the epithelial cells of the thymus which resemble their in vivo counterpart and have thymopoietic ability in reaggregate culture.**

## INTRODUCTION

Over the past two decades, organoids have revolutionized the field of stem cell biology. Recapitulating key elements of the architecture, multicellularity, or function of their native organs on a smaller scale (Rossi et al., 2018), organoids have opened up unprecedented opportunities for personalized medicine. These three-dimensional (3D) structures derived from stem or progenitor cells have been established from a wide variety of organs, particularly of the endodermal lineage (Rossi et al., 2018). However, despite the availability of organotypic cultures (e.g. tissue explants (Anderson and Jenkinson, 1998; Owen and Ritter, 1969) and reaggregates (Giger et al., 2022; Jenkinson et al., 1992; Wagar et al., 2021)) or engineering methods (Kim et al., 2019) (e.g. scaffolds (Asnaghi et al., 2021; Bourgine et al., 2018; Campinoti et al., 2020; Fan et al., 2015; Hun et al., 2016; Poznansky et al., 2000; Purwada and Singh, 2017) and organ-on-a-chip (Goyal et al., 2022)), bona fide immune system-related organoids are considerably underdeveloped. Modelling lymphoid organs is indeed particularly challenging, largely due to the intricate crosstalk between immune and stromal cells required for organ development and function (Anderson and Jenkinson, 1998).

One essential organ for adaptive immunity is the thymus as it functions as the site of T cell development. In the thymus, T cell progenitors undergo lineage commitment and various selection processes to ensure the formation of a diverse, functional, and self-tolerant T cell repertoire, essential for effective immune protection. The instruction of the developing T cells (termed thymocytes) is mostly mediated by thymic epithelial cells (TECs). These stromal cells originate from the pharyngeal endoderm and can be subdivided into cortical and medullary lineages, which mediate successive stages of T cell development.

The essential thymopoietic ability of TECs is however mostly lost in vitro, as traditional two-dimensional (2D) cultures fail to maintain their functionality (Anderson and Jenkinson, 1998; Anderson et al., 1998; Mohtashami and Zúñiga-Pflücker, 2006). Alternative approaches employing OP9 or MS5 cell lines have been developed to circumvent this limitation and study T cell development in vitro (Montel-Hagen et al., 2020; Seet et al., 2017), but the absence of TECs still prevents physiological modelling of T cell selection processes. Other efforts focused on obtaining TECs from pluripotent stem cells (Lai and Jin, 2009; Parent et al., 2013; Ramos et al., 2022; Sun et al., 2013) or through direct reprogramming (Bredenkamp et al., 2014), but these cells largely rely on in vivo grafting to reveal thymopoietic functionality. It was also shown that TECs can form colonies in Matrigel, but these cultures still require feeder cells and their functionality was not demonstrated (Lepletier et al., 2019; Meireles et al., 2017; Wong et al., 2014). Thus, currently the only existing way to preserve TEC functionality in vitro is through (reaggregate) thymic organ cultures, which are organotypic 3D cultures containing different cell types.

Here, in light of what has been achieved for other endoderm-derived epithelia, we postulated that TECs could be grown independently of other cell types as 3D organoids in an extracellular matrix-based hydrogel. We identified culture conditions allowing TECs to form organoids mirroring to some extent the native tissue, and proved their functionality through their ability to mediate T cell development upon reaggregation with T cell progenitors. This work establishes the first thymic epithelial organoids with in vitro thymopoietic ability and is generally the first demonstration of organoids originating from the stromal compartment of a lymphoid organ.

## RESULTS AND DISCUSSION

### Thymic epithelial cells grow and maintain marker expression in defined organoid culture conditions

To establish thymic epithelial organoids, we followed the approach used for other endodermal organs, which included dissociating the tissue, sorting the cells of interest, and seeding them in a basement membrane-rich hydrogel (Matrigel) (Fig. 1A and Fig. S1 A). Since organoids mostly develop from stem or progenitor cells, we focused on the embryonic thymus due to its higher abundance of thymic epithelial progenitor cells compared to the adult organ (Baran-Gale et al., 2020; Kadouri et al., 2020). Although previous attempts to culture TECs often used serum-containing medium (Bonfanti et al., 2010; Campinoti et al., 2020; Wong et al., 2014), we opted for defined organoid basal medium and investigated factors that could promote TEC growth. We hypothesized that mesenchyme-derived factors that have been shown to influence TEC populations both in vivo and in vitro (Alawam et al., 2020; Boehm and Swann, 2013; Chaudhry et al., 2016; James et al., 2021) could also be important for TEC growth in organoid cultures. Among these factors, we found FGF7 of particular interest, as it has recently been shown to sustain the expansion of thymic microenvironments without exhausting the epithelial progenitor pools in vivo (Nusser et al., 2022). Using E16.5 embryonic thymi, we showed that while sorted TECs failed to grow in organoid basal medium, adding FGF7 to the culture supported organoid formation (Fig. 1, B and C). To monitor organoid development, we performed time-lapse imaging from the time of seeding (Fig. 1D and Movie 1) and found that most organoids were derived from single cells with stem/progenitor properties.

**Fig. 1.**
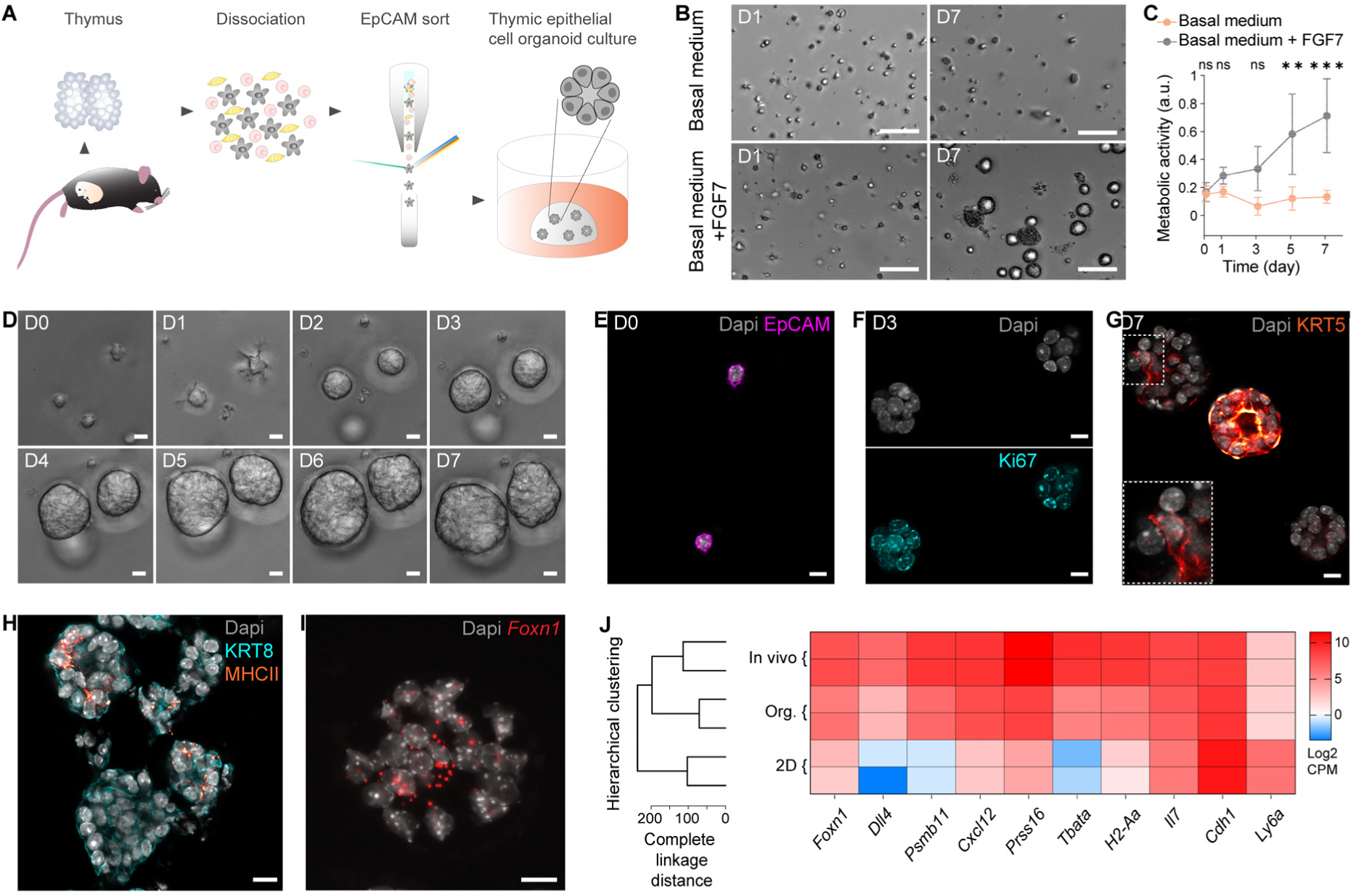
Thymic epithelial cells grow and maintain marker expression in defined organoid culture conditions. (**A**) Schematic of the experimental workflow to isolate, select and culture thymic epithelial cells (TECs) to obtain organoids. (**B**) Brightfield images of TECs in organoid culture conditions one day (D1) and one week (D7) after seeding, in organoid basal medium and in organoid basal medium supplemented with FGF7. Scale bars, 100μm. (**C**) Metabolic activity (measured using resazurin) of TECs cultured in organoid basal medium and in organoid basal medium with FGF7. **: P = 0.0047, ***: P = 0.0005, ns: P > 0.05 (two-way ANOVA; n = 15 per condition, from 3 mice). Data represent mean ± standard deviation (SD). (**D**) One-week time course showing close up brightfield images of sorted TECs from single cells to multicellular organoids. Scale bars, 10μm. (**E**) Immunofluorescence image of individual EpCAM-positive (magenta) TECs immediately after seeding (D0) with nuclei counterstained using Dapi (grey). Scale bar, 10μm. (**F**) Immunofluorescence image of organoids three days after seeding demonstrating that cells undergo proliferation (Ki67 [cyan], Dapi stains nuclei [grey]). Scale bars, 10μm. (**G**) Immunofluorescence image of organoids showing different cell populations after one week in culture, here with medullary cells (KRT5 [red hot]) present in different patterns. Dapi counterstains nuclei (grey). Scale bar, 10μm. (**H**) Immunofluorescence image highlighting MHCII expression (orange) in D7 organoids also stained with KRT8 (cyan) and Dapi (nuclei, grey). Scale bar, 10μm. (**I**) RNAscope image of organoid at D7 showing *Foxn1* expression (red) with nuclei counterstained using Spectral Dapi (grey). Scale bar, 10μm. (**J**) Gene expression profiling. Left: dendrogram (using hclust) showing clustering of thymic epithelial organoids (Org.) with freshly extracted TECs (In vivo) and not TECs cultured in 2D (2D). Metrics is complete linkage distance. Right: heatmap displaying key TEC genes as well as *Cdh1* and *Ly6a* expression for the same three conditions. n = 2 mice per condition.

Immunostaining confirmed that organoids were generated by thymus-derived EpCAM-positive cells (Fig. 1E). Single cells formed small organoids in which a large majority of cells were positive for Ki67 after 3 days (Fig. 1F), and both proliferating and non-proliferating cells were present 4 days later (Fig. S1B). To investigate whether these cell populations could recapitulate TEC diversity, including cortical and medullary types (cTECs and mTECs), we stained organoids for the cTEC marker Keratin 8 (KRT8), as well as for Keratin 5 (KRT5) and with UEA1 lectin as mTEC markers. Overall, our TEC culture system demonstrated a canonical feature of organoids in the emergence of different cell types, with varying degrees of KRT5 expression (Fig. 1G) and the presence of both KRT8-positive and UEA1-reactive populations (Fig. S1C). In addition, at least some organoids were positive for MHCII (Fig. 1H), an important marker of TEC functionality required for the development of CD4+ T cells (Kadouri et al., 2020). TEC differentiation, function and maintenance being critically dependent on the transcription factor *Foxn1* (Žuklys et al., 2016), we further sought to detect transcripts for this master regulator using RNAscope. Unlike in standard 2D cultures where it is highly downregulated (Anderson et al., 1998; Mohtashami and Zúñiga-Pflücker, 2006), a clear *Foxn1* expression could be observed in organoids (Fig. 1I).

To benchmark thymic epithelial organoids against standard 2D culture, we performed bulk RNA sequencing. Unsupervised hierarchical clustering showed the higher transcriptional similarity of thymic epithelial organoids to freshly extracted TECs (in vivo) than to 2D-cultured TECs (Fig. 1J, left). Similarly, a differential expression analysis showed that the expression levels of some key TEC genes, including *Foxn1*, *Dll4* and *Psmb11*, were more similar between in vivo TECs and thymic epithelial organoids compared to 2D-cultured TECs (Fig. 1J, right). Conversely, *Il7* and *Cdh1* were maintained in 2D culture as previously reported (Anderson et al., 1998), and *Ly6a* (a marker of specific TEC subpopulations (Klein et al., 2023)) was upregulated. Lastly, gene set enrichment analysis performed on organoids at different time points confirmed the proliferation peak observed with staining (Fig. S1D).

Collectively, these findings show that the defined culture conditions identified herein allow TECs (i) to grow independently of other cell types (ii) to form organoids containing diverse cell populations and that are transcriptionally similar to in vivo TECs.

### TECs cultured as organoids show in vitro functionality when reaggregated with T cell progenitors

To test the functionality of thymic epithelial organoids (i.e. their ability to mediate T cell development), we recapitulated the well-known reaggregate fetal thymic organ culture (RFTOC) approach, wherein selected thymic cell populations are reaggregated together and cultured at the air-liquid interface (Anderson et al., 1993; Jenkinson et al., 1992). To do so, we dissociated TECs cultured as organoids and reaggregated them with an EpCAM-depleted single cell suspension obtained from E13.5 thymi. We performed EpCAM-depletion in order to keep the mesenchymal cells, which have been proven critical for T cell development (Anderson et al., 1993). We used E13.5 embryonic thymi as source of T cell precursors because they contain thymocytes at the earliest stages of development, prior to the expression of CD4 and CD8 (thus referred to as double negative, DN) (Fig. S2A). This allows to easily monitor whether T cell development happens in the reaggregates. To increase cell number and facilitate handling, mouse embryonic fibroblasts (MEFs) were also added, as done previously (Sheridan et al., 2009). We termed the RFTOCs formed with TECs from the organoid cultures organoid RFTOCs (ORFTOCs) (Fig. 2A).

**Fig. 2.**
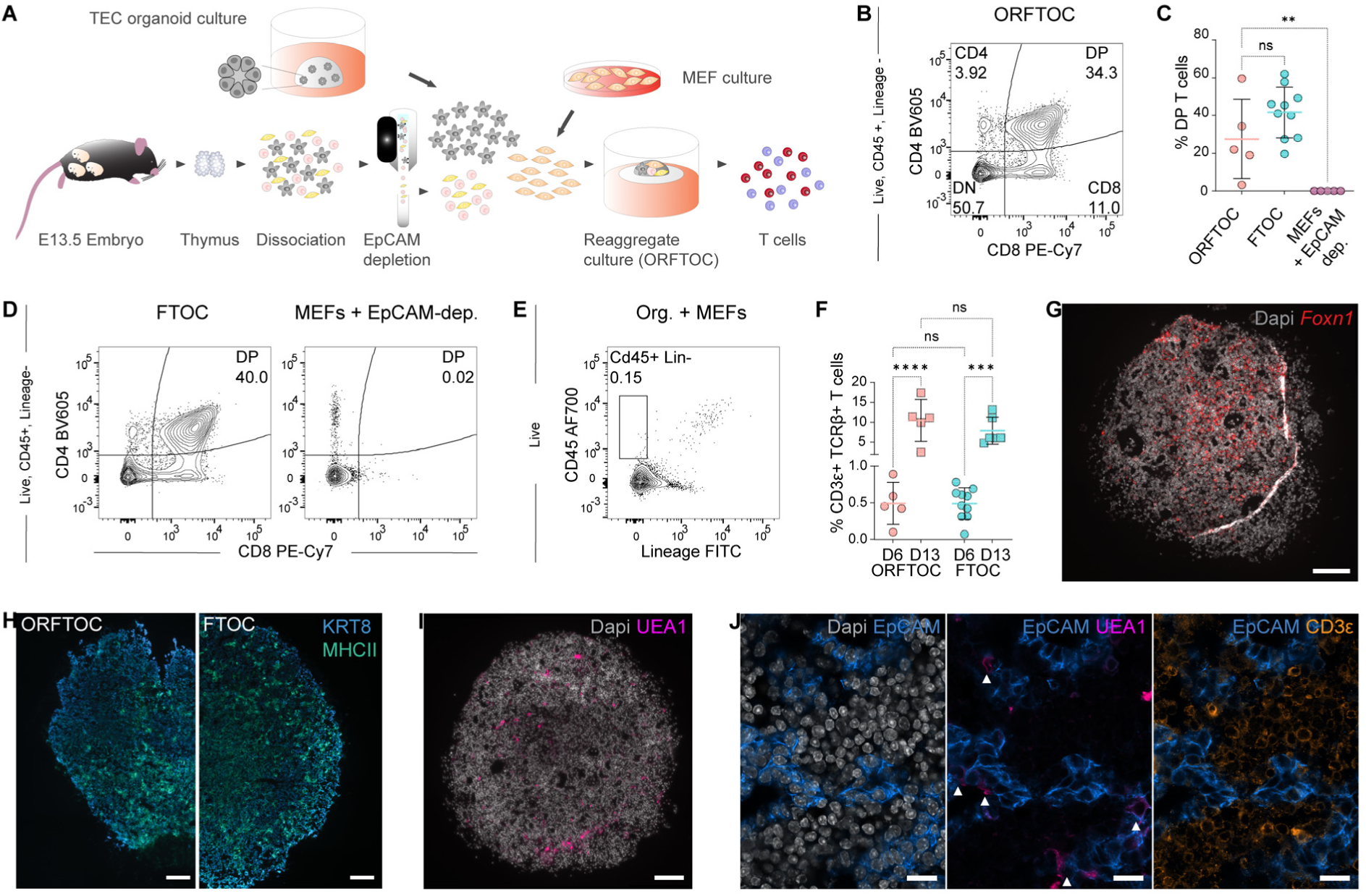
TECs cultured as organoids show in vitro functionality when reaggregated with T cell progenitors. (**A**) Schematic of the experimental workflow to generate Organoid Reaggregate Fetal Thymic Organ Cultures (ORFTOCs) and analyze T cell development. (**B**) Flow cytometry plot showing T cell development in ORFTOCs after 6 days in culture (D6). Gating strategy is indicated on the left. (**C**) Proportion of double positive (DP) thymocytes at D6 in ORFTOCs and controls (fetal thymic organ cultures [FTOCs] and reaggregates containing only mouse embryonic fibroblasts [MEFs] and the EpCAM-depleted cells from thymic lobes). **: P = 0.0081, ns: P > 0.05 (Mood’s median test with P-values adjusted with the false-discovery rate method; n = 5, 10 and 5 for ORFTOC, FTOC and MEFs + EpCAM-depleted cells, respectively, from 5 independent experiments). Graph represents individual datapoints with mean ± SD. (**D**) Flow cytometry plots showing T cell development at D6 in controls (FTOCs [left] and MEFs + EpCAM-depleted cells reaggregates [right]). Gating strategy is indicated on the left. (**E**) Flow cytometry plot showing the absence of a CD45+ Lineage-population in control reaggregates containing only TECs cultured as organoids and MEFs. Gating strategy is indicated on the left. (**F**) Proportion of CD3ε-positive, T cell receptor beta (TCRβ)-positive cells in ORFTOCs and FTOCs at day 6 and 13 (D13). ***: P = 0.0002, ****: P < 0.0001, ns: P > 0.05 (one-way ANOVA with Tukey’s multiple comparisons test; n = 5, 5, 10 and 6 for D6 ORFTOCs, D13 ORFTOCs, D6 FTOCs and D13 FTOCs, respectively, from 5 independent experiments). Graph represents individual datapoints with mean ± SD. (**G**) RNAscope image of D13 ORFTOC section highlighting *Foxn1* expression (red) with nuclei counterstained using Spectral dapi (grey). Scale bar, 100μm. (**H**) Immunofluorescence images of D13 ORFTOC (left) and FTOC (right) sections showing KRT8 (blue) and MHCII staining (green). Scale bar, 100μm. (**I**) Immunofluorescence image of D13 ORFTOC section demonstrating the presence of medullary cells (UEA1-reactivty [magenta]). Dapi counterstains nuclei (grey). Scale bar, 100μm. (**J**) Zoomed immunofluorescence images of D13 ORFTOC section showing epithelial cells (EpCAM [blue]) and nuclei (dapi [grey]) (left), medullary cells (UEA1-reacticity [bright pink]) co-staining with epithelial cells (middle), and T cells (CD3ε [amber]) in-between epithelial cells (right). Scale bars, 100μm.

After 6 days in culture, ORFTOCs were dissociated and analyzed by flow cytometry (Fig. S2B). At this point, thymocytes expressing both CD4 and CD8 (termed double positive, DP) and constituting a developmental stage following the DN phenotype could be readily detected (Fig. 2, B and C), indicating that organoid-derived TECs mediated physiological progression of thymocyte maturation. Notably, the proportion of DPs was similar to that observed in cultured intact thymic lobes (i.e. fetal thymic organ cultures, FTOCs) (Fig. 2, C and D). Conversely, reaggregating only the EpCAM-depleted fraction of E13.5 thymi and MEFs did not yield DP thymocytes (Fig. 2, C and D, and Fig. S2C), demonstrating that organoid-derived TECs are necessary for T cell development in ORFTOCs. Lastly, reaggregates with only organoid-derived TECs and MEFs served as negative control and did not produce immune (CD45+) cells (Fig. 2E), as opposite to the other conditions (Fig. S2B, C, D). To corroborate our findings, we reaggregated organoid-derived TECs with the earliest DN subpopulation (DN1) sorted from adult mice and MEFs, and could also observe thymocyte development (Fig. S2E). The developmental kinetics was however faster in ORFTOCs containing E13.5-derived cells, as expected for first wave early T cell precursors (Rothenberg, 2021).

Extending ORFTOC culture period from 6 to 13 days allowed thymocyte maturation to progress further, as an increased proportion of cells expressed the αβ T cell receptor complex (TCR) (Fig. S2F and Fig. 2F), and differentiated into the separate lineages of CD4+ and CD8+ single positive (SP) thymocyte, respectively (Fig. S2F). FTOCs were again used as reference (Fig. S2G) and demonstrated a comparable proportion of mature SP cells (Fig. S2H).

As expected for functional TECs, ORFTOCs were positive for *Foxn1* (Fig. 2G). Morphologically, ORFTOCs also presented similarities to FTOCs, here highlighted by KRT8 and MHCII staining (Fig. 2H). UEA1 reactivity identified sparse medullary cells throughout ORFTOCs (Fig 2, I and J), and CD3ε staining confirmed the presence of T cells in between EpCAM-positive epithelial cells (Fig. 2J).

In summary, we demonstrated that thymic epithelial organoids maintain their functionality and, when reaggregated with T cell progenitors, mediate T cell development similarly to intact thymic lobe cultures.

### ORFTOCs recapitulate in vivo-like TEC and T cell populations diversity and physiological T cell development

To further characterize the cell types in ORFTOCs, we profiled them and FTOC controls through single cell RNA sequencing (Fig. 3A). This analysis revealed three main clusters corresponding to the epithelial, immune, and mesenchymal compartments of ORFTOCs (Fig 3B). Unsupervised clustering identified 7 main clusters of epithelial cells (Fig. S3A), which we annotated according to in vivo datasets (Baran-Gale et al., 2020; Bautista et al., 2021; Bornstein et al., 2018; Gao et al., 2022; Nusser et al., 2022; Park et al., 2020): ‘early cTECs’, ‘cTECs’, ‘early mTECs’, ‘pre-Aire mTECs’, ‘Aire and Spink5 mTECs’, ‘tuft-like mTECs’, and ‘adult bipotent progenitor-like’ (Fig. S3B). For the immune cells, clusters covered the main T cell developmental stages defined in vivo (Cordes et al., 2022; Luis et al., 2016; Mingueneau et al., 2013; Park et al., 2020; Rothenberg, 2021; Zhou et al., 2019), spanning from progenitors to mature T cells (Fig. S3, C and D). Both ORFTOCs and FTOCs contributed to all subpopulations (Fig. 3, C and D), suggesting that ORFTOCs faithfully recapitulate the different cell types present in FTOC controls. The biggest differences were observed for clusters representing cTECs and early stages of T cell development: (i) more ‘early cTECs’ were present in FTOCs and (ii) more ‘cTECs’ and ‘thymus-seeding progenitor (TSP) to DN early 1’ and ‘TSP to DN early 2’ cells were present in ORFTOCs. A potential explanation for this is the difference in embryonic age between the epithelial cells used for organoids and FTOCs, as FTOCs were from E13.5 thymi to match T cell progenitors in ORFTOCs, while TECs in ORFTOCs were from E16.5 thymi.

**Fig. 3.**
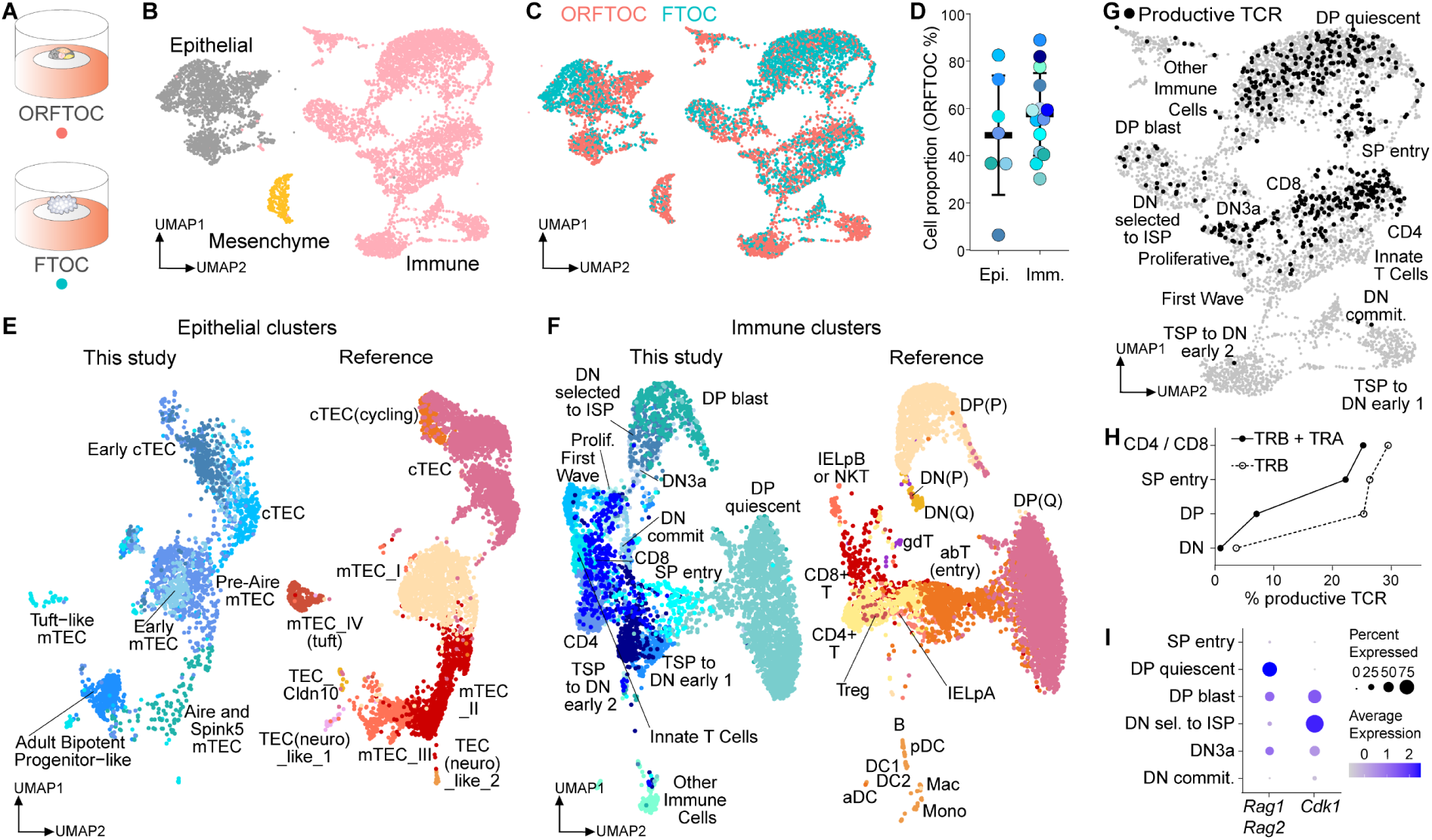
ORFTOCs recapitulate in vivo-like TEC and T cell populations diversity and physiological T cell development. (**A**) Schematic of the conditions used for ORFTOC and FTOC single-cell RNA sequencing with hashtag antibodies (HTOs) and analyzed after 13 days in culture. (**B**) Uniform Manifold Approximation and Projection (UMAP) showing 3 main clusters corresponding to the main input populations (epithelial, immune and mesenchymal cells). (**C**) UMAP displaying ORFTOC and FTOC cells distribution in the different clusters. (**D**) Dot plot representing ORFTOC proportion for each cluster (dot) within the epithelial or immune main populations, as well as mean ORFTOC proportion and standard deviation. Dot colors are matching clusters colors (Fig. S3, A and C). No outliers within epithelial or immune compartments were identified by Grubbs test. (**E - F**) UMAPs showing the integration of the epithelial (E) and immune (F) clusters identified in this study (left) with the mouse dataset of the reference atlas by Park et al. (Park et al., 2020) (right). (**G**) UMAP of the immune cluster (grey), highlighting cells identified as productive and bearing both TCR chains (black). (**H**) Proportion of productive cells with rearranged TRB or both TRA and TRB chains for the main thymocyte developmental stages. (**I**) Dot plot representing the average expression level and the percentage of cells expressing the recombination enzymes *Rag1* and *Rag2* as well as the cyclin protein *Cdk1* during the recombination and proliferation stages of thymocyte development. Epi: epithelial, Imm: immune, cTEC: cortical TECs, mTEC: medullary TECs, DN: double negative, ISP: intermediate single positive, Prolif: proliferative, commit: commitment, TSP: thymus-seeding progenitors, DP: double positive, (P): proliferative, (Q): quiescent, IELpB: intestinal intraepithelial lymphocytes precursor B, NKT: natural killer T, IELpA: intestinal intraepithelial lymphocytes precursor A, DC: dendritic cells, pDC: plasmacytoid dendritic cells, aDC; activated dendritic cells, Mac: macrophages Mono: monocytes, sel: selected.

To compare our in vitro populations with the in vivo thymus, we aligned our clusters to the mouse dataset of the reference atlas by Park et al. (Park et al., 2020) (Fig. 3, E and F). We found strong overlap in most epithelial cell types (Fig. 3E), with the cTECs aligning together and most in vitro mTEC clusters matching their in vivo counterparts. However, the ‘adult bipotent progenitor-like’ cluster was smaller in vivo compared to in vitro. Immune clusters from our dataset also matched clusters defined for in vivo populations (Fig. 3F), especially from the ‘DP blast’/‘DP (P)’ stage onwards and, most importantly, for the CD4 and CD8 stages (mature T cells).

Besides gene expression, we also studied in vitro TCR recombination dynamics through V(D)J sequencing, allowing us to map productive T cells bearing both TCR chains on the immune UMAP (Fig. 3G). The quantification of productive chains presenting all V(D)J regions showed that the recombination of the TCRβ (TRB) and -α (TRA) chains were mostly achieved prior to and at the DP stage, respectively (Fig. 3H), similarly to the Park dataset (Park et al., 2020). In addition, thymocytes underwent proliferation (marked by high *Cdk1* expression) in between the recombination stages (marked by high *Rag1* and *Rag2* expression) (Fig. 3I), which also aligns with in vivo data (Park et al., 2020; Rothenberg, 2021).

Taken together, these results show the transcriptional similarity of ORFTOCs to FTOCs and that ORFTOCs preserve in vivo-like TEC diversity and T cell development.

### ORFTOCs show thymus-like ability to attract new T cell progenitors and improved epithelial organization upon in vivo grafting

The thymus continuously attracts bone marrow-derived hematopoietic precursors and commits them to the T cell lineage (Lai and Kondo, 2007; Lavaert et al., 2020). To test whether ORFTOCs retain this crucial capacity, we transplanted them under the kidney capsule of syngeneic CD45.1 recipient mice (Fig. 4A). After 5 weeks, all grafts developed into sizeable thymus-like tissues (4/4 ORFTOCs [Fig. 4B], 4/4 FTOC controls). Using flow cytometric analyses (Fig. S4A), we identified all major thymocyte populations (DN, DP, CD4, CD8) in ORFTOC grafts (Fig. 4C), and their proportions were comparable to FTOC control grafts and control thymi (Fig. 4D). This result demonstrated that normal αβ-TCR T cell lineage maturation was supported in ORFTOC grafts. A further detailed analysis (Fig. S4, B and C) detected thymocytes at the DN3 to DN4 transition at the time of ORFTOC graft retrieval, a stage attesting to successful β-selection (Rothenberg, 2021). In addition, DP thymocytes expressing CD69, which indicates positive selection-induced TCR signaling (Steier et al., 2023), were present (Fig. S3 B and D). Together, these results illustrate ORFTOC graft ability to continuously attract and select blood-borne T cell progenitors. Finally, the presence of CD45.1+ mature thymocytes (classified as M1 [CD24+ CD69+] and M2 [CD24+ CD69-], respectively [Fig. 4E and Fig. S4E]) and of regulatory T cells (Fig. 4F and Fig. S4F) showed ORFTOC graft capacity to generate mature CD4 and CD8 T cells, impose their post-selection maturation and select T cells with a regulatory phenotype.

**Fig. 4.**
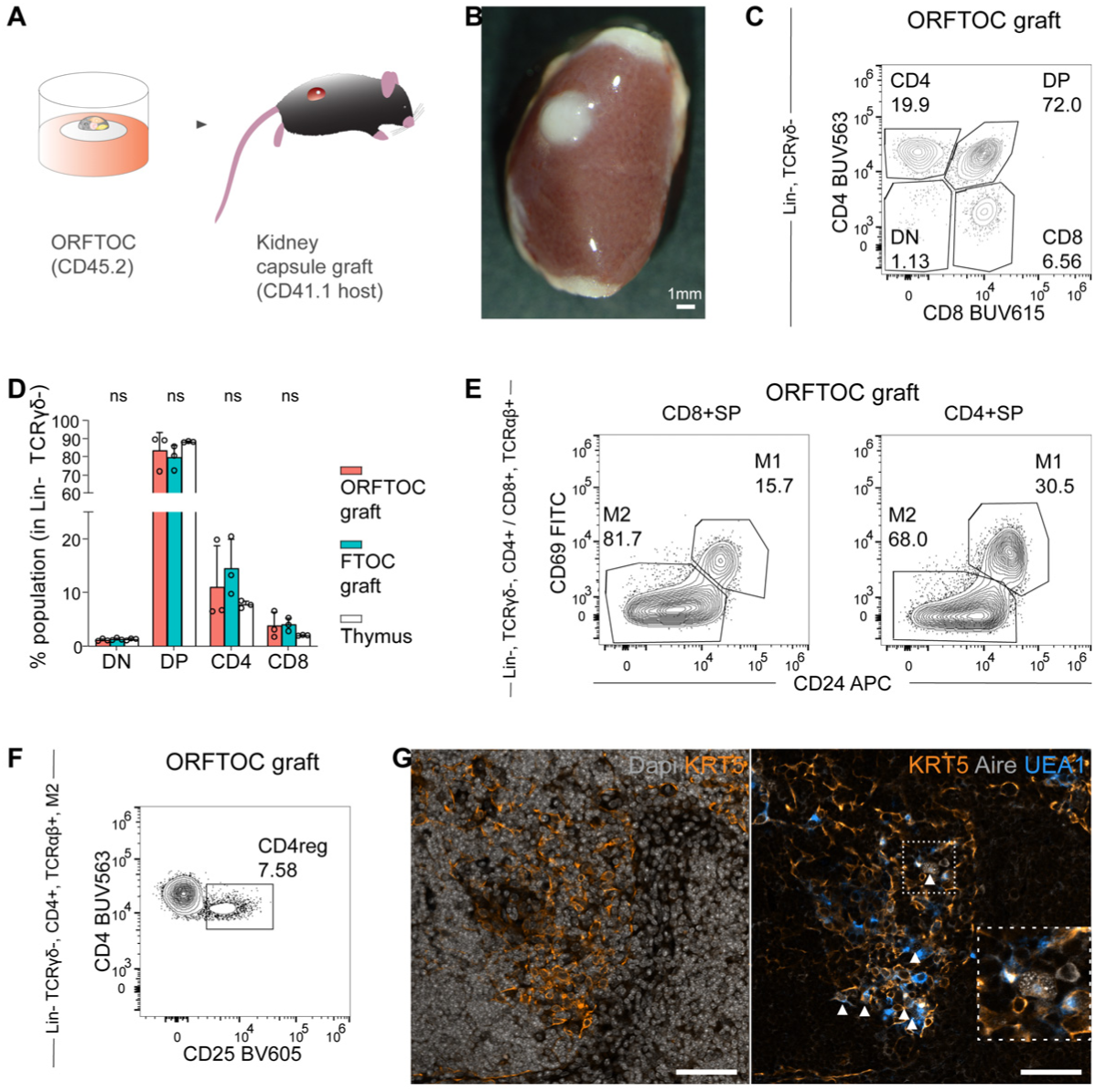
ORFTOCs show thymus-like ability to attract new T cell progenitors and improved epithelial organization upon in vivo grafting. (**A**) Schematic representing the experimental design for the grafting of ORFTOCs under the kidney capsule. (**B**) Widefield image of an ORFTOC graft retrieved after 5 weeks. Scale bar, 1mm. (**C**) Flow cytometry plot showing host thymocyte development in ORFTOC grafts. Gating strategy is indicated on the left. (**D**) Proportion of the major thymocyte subpopulations in ORFTOC grafts, in control FTOC grafts and thymi. ns: P > 0.05 (one-way ANOVA for each subpopulation between conditions; n = 3 grafts/mice for each condition). Bar graph represents mean ± SD and individual datapoints. (**E**) Flow cytometry plots showing two separate post-selection stages (M1 and M2) within the CD8+ and CD4+ SP populations in ORFTOC grafts. Gating strategy is indicated on the left. (**F**) Flow cytometry plot highlighting the presence of CD4 regulatory T cells (CD4reg) within the M2 population in ORFTOC grafts. Gating strategy is indicated on the left. (**G**) Immunofluorescence images of ORFTOC graft section. Left: medullary cells (KRT5 [amber]) are present in the less dense area (Dapi [grey]). Right: UEA1-reactive (azure) and Aire-positive cells (grey, highlighted with arrowhead) are also present in the medullary region. Scale bars, 100μm.

Histological staining showed that ORFTOC grafts, similar to FTOC grafts, displayed the characteristic differences in cellular densities between cortical and medullary areas seen in the native thymus (Gordon and Manley, 2011) (Fig. S4G and Fig. 4G). Immunostaining confirmed the presence of medullary areas (positive for KRT5 and reactive to UEA1) containing Aire-positive cells (Fig. 4G). These medullary areas were larger, better organized and more mature compared to those observed after in vitro culture only (Fig. 2I), likely due to continuous seeding with new T cell progenitors and prolonged crosstalk with immune cells (Irla et al., 2010).

In conclusion, kidney capsule transplants showed that organoid-derived TECs in ORFTOCs have the (i) capacity to mature and reach an organization resembling the native thymus and (ii) long-term ability to attract T cell progenitors and mediate physiological T cell development.

In this study, we showed that stromal cells of a lymphoid organ, namely epithelial cells of the thymus, can be cultured as organoids similarly to cells from other endoderm-derived organs. We established TEC-specific culture conditions, characterized the organoids, and demonstrated their superiority in maintaining TEC marker expression compared to conventional 2D cultures. Reaggregating TECs from organoid cultures with T cell progenitors proved their functionality and ability to mediate T cell development. TEC and T cell populations in reaggregates resembled the native cell types, and T cell maturation was recapitulated in a physiological manner. Finally, kidney capsule transplants demonstrated the long-term capability of organoid reaggregates to attract new T cell progenitors and mediate their entire development.

Overall, this work addressed a long-standing challenge in the thymus field and presents the first method to culture TECs independently of other cell types in a way that maintains their thymopoietic ability. Although thymic epithelial organoids recapitulate many key organoid features such as cell population diversity and possibility to be expanded and passaged, maintaining their functionality in the long term remains challenging. This is probably linked to some niche factors missing in the current relatively minimal culture conditions, which over time either generally prevent functionality to be maintained or enrich for specific subsets that might lack functionality (Gao et al., 2022). Future work including single-cell transcriptomic analysis of the organoids will most likely help identify yet unexplored but necessary niche factors.

Thymic epithelial organoids nevertheless open up new opportunities to study T cell development in vitro in a physiological manner and gain new insights into TEC biology. As TECs undergo deterioration during aging and different medical conditions, the development of the current and future culture conditions might also pave the way for novel thymus regeneration strategies. Finally, to the best of our knowledge, the generation of bona fide organoids from the stromal compartment of a lymphoid organ is unprecedented.

## Materials and Methods

### Mice

For all in vitro experiments, C57BL/6J mice were purchased from Charles River France and maintained in the EPFL animal facility until use. For grafting experiments, Ly5.1 and C57BL/6J mice were bred and maintained in the mouse facility of the Department of Biomedicine at the University of Basel. For timed mating, noon of the day of the vaginal plug was considered as day 0.5 of embryonic development (E0.5). Mice were housed in individual cages at 23°C ± 1°C with a 12 h light/dark cycle, and supplied with food and water ad libitum. All animal work was conducted in accordance with Swiss national guidelines, reviewed and approved by the Cantonal Veterinary Offices of Vaud and of Basel-Stadt, license numbers VD3035.1, VD3823 and BS2321.

### Isolation of thymic epithelial cells (TECs)

E16.5 embryonic thymi were dissected and collected in Eppendorf tubes containing FACS buffer (PBS [Gibco Catalog No. 10010-015] + 2% fetal bovine serum [FBS] [Thermo Fisher Scientific, Catalog No. 26140079]). Lobes were rinsed with PBS and digested with 475 µl TrypLE (Gibco Catalog No. 12605-028) for 5 min at 37 °C under agitation (Eppendorf, ThermoMixer C). Lobes were pipetted to promote dissociation, 25 μl DNase (Sigma-Aldrich, Catalog No. 10104159001, from 1 mg/ml stock) was added and the tubes were incubated for another 5 min. Lobes were again pipetted to help dissociation, TrypLE was quenched with 1ml Adv. DMEM/F12 (Thermo Fisher Scientific, Catalog No. 12634028) containing 10 % FBS and the cell suspension was filtered through a 40 μm strainer. The cells were pelleted and resuspended in FACS buffer for staining with the following antibodies for 20 min at 4° C: Ter119-FITC (BioLegend, Catalog No. 116205, 1/100), EpCAM-PE (BioLegend, Catalog No. 1198206, 1/80), PDGFR-α-APC (BioLegend, Catalog No. 135907, 1/40), PDGFR-β-ACP (BioLegend, Catalog No. 136007, 1/40), CD31-PE-Cy7 (BioLegend, Catalog No. 102524, 1/160), MHCII-APC/Fire750 (BioLegend, Catalog No. 107651, 1/160), CD45-Pacific Blue (BioLegend, Catalog No. 103126, 1/200). Dapi (Tocris, Catalog No. 4748, 0.5 ug/ml) was used to exclude dead cells. After staining, the antibodies were washed and the cells resuspended in FACS buffer for sorting using an Aria Fusion (BD). The sorting strategy for isolating thymic epithelial cells is shown in Fig. S1 A. Sorted cells were collected in TEC medium (see below) containing 2 % FBS and 2.5 μM Thiazovivin (Stemgen, Catalog No. AMS.04-0017).

### Thymic epithelial organoid culture

Sorted thymic epithelial cells were embedded in growth–factor-reduced Matrigel (Corning, Catalog No. 356231) (∼1.55 × 10^4^ cells per 20 mL drop) and plated in 24-well plates (Corning, Catalog No. 353047, or Ibidi, Catalog No. 82426). After Matrigel polymerization, TEC medium was added. TEC medium consisted of organoid basal medium (Advanced DMEM/F-12 supplemented with 1× GlutaMAX [Thermo Fisher Scientific, Catalog No. 35050038], 10 mM HEPES [Thermo Fisher Scientific, Catalog No. 15630056], 100 μg ml^−1^ Penicillin–Streptomycin [Thermo Fisher Scientific, Catalog No. 15140122], 1× B-27 supplement [Thermo Fisher Scientific, Catalog No. 17504001], 1× N2 supplement [Thermo Fisher Scientific, Catalog No. 17502001], 1 mM N-Acetylcysteine [Sigma-Aldrich, Catalog No. A9165]) plus 100 ng ml^−1^ FGF7 (Peprotech, Catalog No. 100-19). 2.5μM Thiazovivin was also added to the medium for the first two days. Medium was changed every second day. Organoids were cultured at 37°C with 5% CO_2_.

### Organoid proliferation assays

Sorted thymic epithelial cells were embedded in 10 μl Matrigel drops (∼7.5 x 10^3^ cells/drop) in a 48-well plate (Corning, Catalog No. 353078). On the day of seeding (day 0), at day 1, 3, 5 and 7, 220 μM resazurin (Sigma-Aldrich, Catalog No. R7017) was added to organoid basal medium and incubated with the cells for 4 h at 37 °C. Afterwards, the resazurin-containing medium was collected and replaced by fresh TEC medium with or without FGF7. Organoid proliferation was estimated by measuring the reduction of resazurin to fluorescent resorufin in the medium using a Tecan Infinite F500 microplate reader (Tecan) with 560 nm excitation and 590 nm emission filters. For analysis, data were normalized from minimum to maximum.

### Bulk transcriptome profiling

Sorted thymic epithelial cells were culture as indicated above. As controls, sorted thymic epithelial cells from E16.5 embryos were either directly lysed in RLT buffer (QIAGEN, Catalog No. 74004) containing 40 mM DTT (ITW Reagents, Catalog No. A2948) or cultured in 2D on plates coated with 6 μg/ml laminin (R&D Systems, Catalog No. 3446-005-01). Cultures were done in TEC medium. Organoids were collected in cold PBS to dissolve Matrigel and then lysed in RLT buffer with DTT. They were collected after 3 and 7 days. Cells cultured in 2D were directly collected in RLT buffer with DTT. They were collected once a confluent monolayer formed, after 3 days, as prolonged culture in these conditions lead to cell detachment and death. RNA was extracted using QIAGEN RNeasy Micro Kit (QIAGEN, Catalog No. 74004) according to manufacturer’s instructions. Purified RNA was quality checked using a TapeStation 4200 (Agilent), and 88 ng were used for QuantSeq 3′ mRNA-Seq library construction according to manufacturer’s instructions (Lexogen, Catalog No. 015.96). Libraries were quality checked using a Fragment Analyzer (Agilent) and were sequenced in a NextSeq 500 (Illumina) using NextSeq v2.5 chemistry with Illumina protocol #15048776. Reads were aligned to the mouse genome (GRCm39) using star (version 2.7.0e). R (version 4.1.2) was used to perform differential expression analyses. Count values were imported and processed using edgeR (Robinson et al., 2010). Expression values were normalized using the trimmed mean of M values (TMM) method and lowly-expressed genes (< 1 counts per million) and genes present in less than three samples were filtered out. Differentially expressed genes were identified using linear models (Limma-Voom) (Smyth et al., 2018), and P-values were adjusted for multiple comparisons by applying the Benjamini-Hochberg correction method (Reiner et al., 2003). Voom expression values were used for hierarchical clustering using the function hclust (Murtagh, 1987) with default parameters, and for heatmap generation. Single sample gene set enrichment analysis (GSEA) (Subramanian et al., 2005) was used to score the E2F targets hallmark proliferation gene set (Howe et al., 2018; Liberzon et al., 2015) between samples.

### Whole-mount immunofluorescence staining

Organoid samples were fixed in 4% paraformaldehyde (Thermo Fisher Scientific, Catalog No. 15434389) in PBS for 30 min at room temperature and subsequently washed with PBS. Samples were permeabilized in 0.2 % Triton X-100 (Sigma-Aldrich, Catalog No. T8787), 0.3 M glycine (Invitrogen, Catalog No. 15527-013) in PBS for 30 min at room temperature and blocked in 10 % serum (goat [Thermo Fisher Scientific, Catalog No. 16210064] or donkey [Abcam, Catalog No. ab7475]), 0.01% Triton X100 and 0.3M glycine in PBS for 4h at room temperature. Samples were then incubated with primary antibodies overnight at 4 °C, washed with PBS, incubated with secondary antibodies overnight at 4 °C, and washed with PBS. Mounting was done with Fluoromount-G (SouthernBiotech, Catalog No. 0100-01). The following primary and secondary antibodies were used: MHCII-Biotin (BioLegend, Catalog No. 107603, 1/200), UEA1 (Vector Laboratories Catalog No. B-1065, 1/500), Keratin 5 (BioLegend, Catalog No. 905501, 1/200), Keratin 8 (Abcam Catalog. No. ab53280, 1/200), Ki67 (BD Pharmingen, Catalog No. 550609, 1/200), EpCAM-APC (Invitrogen, Catalog No. 17-5791-82, 1/200), Streptavidin Alexa 488 (Thermo Fisher Scientific, Catalog No. S-11223, 1/500), Streptavidin Alexa 647 (Thermo Fisher Scientific, Catalog No. S-21374, 1/500), Goat anti-Rat Alexa 647 (Thermo Fisher Scientific, Catalog No. A-21247, 1/500), Donkey anti-Mouse Alexa 568 (Thermo Fisher Scientific, Catalog No. A-10037, 1/500), Donkey anti-Rabbit Alexa 488 (Thermo Fisher Scientific, Catalog No. A-21206, 1/500), Donkey anti-Rabbit Alexa 568 (Thermo Fisher Scientific, Catalog No. A-11077, 1/500). Dapi (Tocris, Catalog No. 47481 mg/ml) was used to stain nuclei.

### Reaggregate culture

E13.5 embryonic thymi were dissected and collected in Eppendorf tubes containing FACS buffer. Lobes were rinsed with PBS and digested with 475 ul TrypLE and 25 μl DNase (from 1 mg/ml stock) for 5 min under agitation. Lobes were pipetted to help dissociation and TrypLE was quenched with 1ml Adv. DMEM/F12 containing 10 % FBS. The cells were pelleted and resuspended in FACS buffer for immunomagnetic cell separation with EpCAM-conjugated beads (Miltenyi Biotec, Catalog No. 130-105-958, 1/4). After 20 min incubation at 4°C, the unbound complexes were washed and the cells processed through magnetic columns (Miltenyi Biotec, Catalog No. 130-042-401) following manufacturer instruction. The EpCAM-depleted fraction was collected and used to prepare reaggregates with dissociated thymic epithelial organoids and mouse embryonic fibroblasts (MEFs).

Thymic epithelial organoids at day 7 of culture were collected in cold Advanced DMEM/F-12 supplemented with 1× GlutaMAX, 10 mM HEPES and 100 μg ml^−1^ Penicillin–Streptomycin. Organoids were pelleted and digested with 950 μl TrypLE and 50 μl DNase (from 1 mg/ml) for 5min at 37 °C. Organoids were pipetted to improve dissociation. In case digestion was insufficient, organoids were further digested for 5 min with Trypsin + 0.25% EDTA (Gibco, Catalog No. 25200-072) at 37 °C and pipetted until the obtention of a single cell suspension. Dissociation was quenched with Adv. DMEM/F12 containing 10 % FBS and the cells pelleted.

Wild-type MEFs were a kind gift from the Blackburn laboratory. MEFs were cultured in Advanced DMEM/F-12 supplemented with 1× GlutaMAX, 1x Non-Essential Amino Acids (Gibco, Catalog No. 11140035), 100 μg ml^−1^ Penicillin–Streptomycin and 10 % FBS on gelatin-coated dishes (0.1% gelatin in H_2_O) (Sigma-Aldrich, Catalog No. G1890). MEFs were harvested using Trypsin EDTA 0.25% for 2 min at 37 °C. Dissociation was quenched with Adv. DMEM/F12 containing 10 % FBS and the cells pelleted.

For reaggregates using adult double negative 1 (DN1) thymocytes as input population, adult thymi were dissected from 4 weeks old female C57BL/6J mice. Thymi were cut in small pieces with a scalpel to liberate thymocytes, which were filtered to a single cell suspension with a 40 μm strainer. Cells were incubated with APC anti-mouse CD8a Antibody (BioLegend, Catalog No. 100711, 1/50) for 20 min at 4°C in FACS buffer and washed. Cells were then incubated with anti-APC magnetic beads (Miltenyi Biotec, Catalog No. 130-090-855, 1/4) for 20 min at 4°C. The unbound beads were washed away and the cells processed through magnetic columns following manufacturer instruction. The APC depleted fraction was collected and used for staining with the following antibodies: Ter119-FITC (BioLegend, Catalog No. 116205, 1/800), Cd45R-FITC (BioLegend, Catalog No. 103205, 1/800), CD11b-FITC (Thermo Fisher Scientific, Catalog No. 11-0112-82, 1/800), Ly-6G-FITC (BioLegend, Catalog No. 108405, 1/800), Cd11C-FITC (BioLegend, Catalog No. 117306, 1/800), NK-1.1-FITC (BioLegend, Catalog No. 108705, 1/800), CD3-FITC (BioLegend, Catalog No. 100306), CD4-FITC (BioLegend, Catalog No. 100510), CD45-Pacific Blue (BioLegend, Catalog No. 103126, 1/200) or CD45-AF700 (BioLegend, Catalog No. 103127, 1/160), CD44-PE (BioLegend, Catalog No. 103008, 1/160), CD25-BV711 (BioLegend, Catalog No. 102049, 1/160) and Dapi (Tocris, Catalog No. 4748, 0.5 ug/ml). After staining, the antibodies were washed and the cells resuspended in FACS buffer for sorting using an Aria Fusion (BD). The sorting strategy for isolating DN1 thymocytes was gating on cells, single cells, live cells, CD45+ cells, CD44+ CD25-cells. DN1 thymocytes were collected in ORFTOC medium (see below).

Organoids reaggregate fetal thymic organ culture (ORFTOCs) were prepared as previously described (Sheridan et al., 2009). Briefly, the cell suspension for each ORFTOC typically contained 10^5^ EpCAM-depleted cells, 10^5^ thymic epithelial organoid cells, and 10^5^ MEFs (or 10^5^ thymic epithelial organoid cells, 4×10^4^ DN1 thymocytes and 10^5^ MEFs). These cells were transferred to an Eppendorf tube and pelleted. The pellet was resuspended in 60 μl of the medium used for culture, and transferred to a tip sealed with parafilm inside a 15 ml Falcon tube. Cells were pelleted inside the tip for 5 min at 470 rcf. The pellet was then gently extruded on top of a filter membrane (Merck, Catalog No. ATTP01300) floating on culture medium in 24well plate. ORFTOC culture medium consisted of advanced DMEM/F-12 supplemented with 1× GlutaMAX, 1x Non-Essential Amino Acids, 100 μg ml^−1^ Penicillin–Streptomycin, 2 % FBS and 100 ng/ml FGF7. 2.5μM Thiazovivin was added for the first two days of culture and half of the medium volume was changed every second day.

Controls where one of the cell population is absent were made the same way. For FTOC controls, E13.5 dissected lobes were directly placed on top of a filter membrane and also cultured in ORFTOC medium.

All cultures were done at 37°C with 5% CO_2_.

### Flow cytometry analysis of ORFTOCs and FTOCs

After 6 and 13 days in culture, ORFTOCs, FTOCs, reaggregates with DN1 thymocytes and controls reaggregates were gently detached from the filter membrane by pipetting and transferred to Eppendorf tubes, together with the culture medium to collect recently emigrated T cells. Samples were pelleted, rinsed with PBS and digested with 200 μl TrypLE for 10 min at 37° C with agitation on an Eppendorf shaker (800 rpm). Dissociation was quenched with 1ml Adv. DMEM/F12 containing 10 % FBS and the cells pelleted. Cells were resuspended in FACS buffer for staining. The cells were incubated for 20 min with the following antibodies: Ter119-FITC (BioLegend, Catalog No. 116205, 1/800), Cd45R-FITC (BioLegend, Catalog No. 103205, 1/800), CD11b-FITC (Thermo Fisher Scientific, Catalog No. 11-0112-82, 1/800), Ly-6G-FITC (BioLegend, Catalog No. 108405, 1/800), Cd11C-FITC (BioLegend, Catalog No. 117306, 1/800), NK-1.1-FITC (BioLegend, Catalog No. 108705, 1/800) together referred as Lineage, CD44-PE (BioLegend, Catalog No. 103008, 1/160), CD69-APC (BioLegend, Catalog No. 104513, 1/160), CD4-BV605 (BioLegend, Catalog No. 100548, 1/40), CD3-PerCP/Cy5.5 (BioLegend, Catalog No. 100327, 1/160), CD8-PE/Cy7 (BioLegend, Catalog No. 100722, 1/160), CD25-BV711 (BioLegend, Catalog No. 102049, 1/160), CD45-AF700 (BioLegend, Catalog No. 103127, 1/160), TCRβ-BV421 (BioLegend, Catalog No. 109230, 1/80) and Dapi (Tocris, Catalog No. 4748, 0.5 ug/ml). After staining, the antibodies were washed and the cells resuspended in FACS buffer for analyzing using a LSR Fortessa Cytometer (BD). The gating strategy for analysis is shown in Fig. S2 B. Beads (UltraComp, Thermo Fisher Scientific Catalog No. 01-3333-42) were used for single color staining for compensation. Gates were based on T cells extracted from a young adult. Flow cytometry data were analyzed using FlowJo (BD, version 10.9.0).

### Single-cell transcriptome profiling

After 13 days in culture, ORFTOC and FTOC samples were collected and dissociated as described for flow cytometry analysis. After dissociation, two ORFTOC samples and two FTOC samples were pooled, respectively. For each pool, 500 000 cells were incubated with 1ul TotalSeq Antibody (HTO) (BioLegend, Catalog No. 155863 and 155861) in 50 μl FACS buffer for 30 min on ice. Antibodies were washed two times with FACS buffer and the single cell suspensions filtered through a 40 μm strainer. After cell count, samples were mixed in a 1:1 ratio and processed using Chromium Next GEM Single Cell 5’ Reagent Kits v2 (Dual Index) with Feature Barcode technology for Cell Surface Protein & Immune Receptor Mapping reagents (10X Genomics, Catalog No. PN-1000265, PN-1000256, PN-1000190, PN-1000287, PN-1000215 and PN-100025) following manufacturer’s instruction. Single Cell Mouse TCR amplification Kit (10X Genomics Catalog No. 1000254) was used to prepare TCR libraries. Sequencing was done using NovaSeq v1.5 STD (Illumina protocol #1000000106351 v03) for around 100,000 reads per cell. The reads were aligned using Cell Ranger v6.1.2 to the mouse genome (mm10). Raw count matrices were imported into R and analyzed using Seurat v4.2.0 (Hao et al., 2021). HTO with less than 100 features and less than 1 count were discarded. Cells with less than 600 features, less than 0.4 or more than 10 percent mitochondrial genes were discarded. Demultiplexing was performed using HTODemux with standard parameters. Doublets were removed using recoverDoublets from scDblFinder package (Germain et al., 2022) and based on doublets identified from HTOs. Data were normalized using SCTransform and with cell cycle score as variable to regress. The three clusters representing the main cell types were obtained using PCA and UMAP with 18 dimensions and a resolution of 0.005. Each cell type was then subset and thresholded based on *EpCAM*, *Ptprc* and *Pdgfra* expression. Epithelial clusters were identified using 18 dimensions and a resolution of 0.4, leading to 7 clusters that were named based on markers from previous datasets (Baran-Gale et al., 2020; Bautista et al., 2021; Bornstein et al., 2018; Gao et al., 2022; Kernfeld et al., 2018; Park et al., 2020). Immune clusters were identified using 18 dimensions and a resolution of 3. Immune clusters were further merged to obtain 14 clusters representing main T cell developmental stages based on markers from previous datasets (Cordes et al., 2022; Mingueneau et al., 2013; Park et al., 2020; Rothenberg, 2021). The number of cells per clusters in both FTOC and RFTOC samples were calculated to show HTO repartition between both samples. TCR analysis was conducted using scRepertoire (Borcherding et al., 2020). Filtered contig output from Cell Ranger was used as input and added to immune cells metadata. Productive cells with both TRA and TRB chains were plotted on the UMAP, and percentage of productive cell (either at least TRB chain with no NA and no double chain, or both TRA and TRB chains with no NA and double chain accepted only for TRA) per cluster calculated. The mouse samples from the dataset from Park et al. (Park et al., 2020) were used for alignment. H5ad files were converted to Seurat object, TECs were subset from the stromal dataset and 4 weeks-old T cells from the mouse total dataset. Alignment was performed using SCTransform and canonical correlation analysis (CCA) with the Park dataset labeled as reference and otherwise default parameters.

### Kidney capsule grafting and analysis

ORFTOCs were grafted in CD45.1 host mice and FTOC controls in CD45.2 host mice. Mice were treated with the analgesic Carprofen (10 mg/kg in drinking water) 12-24 h prior to transplantation. Mice were anesthetized with Ketalar/Rompun (100 mg/kg Ketamin and 20 mg/kg Xylazin, intraperitoneal). Lacrinorm eye gel (Bausch & Lomb) was administered to avoid dehydration of the cornea during the procedure. Anesthetized mice were shaved laterally and disinfected using Betadine. The surgery was performed on a heating pad in order to minimize body temperature drop. A small incision of approximately 1 cm was done first on the skin and then in the peritoneum. By pulling at the posterior fat of the kidney with forceps, the kidney was exposed outside of the peritoneum and kept wet with PBS. Under the microscope, an incision and a channel were done with watchmaker-forceps on the kidney capsule’s membrane and one ORFTOC or FTOC was placed under the membrane. After positioning the kidney back into the peritoneum, the wound was closed with two stitches (resorbable suture material 5/0; Polyactin 910; RB-1 plus; Johnson&Johnson). The skin opening was closed with staples, which were removed 7-10 days later. An analgesic (Temgesic, Buprenorphine 0.1 mg/kg, subcutaneous) was administered at the end of the procedure followed by continuous treatment of transplanted mice by Carprofen (10 mg/kg in drinking water) for 3 days. After the transplants, mice were monitored daily and weighed every second day to confirm their wellbeing. Grafts were analyzed 5 weeks after transplantation.

At the time of analysis, mice were sacrificed with CO_2_ and kidneys retrieved. Grafts were separated from the kidney under the microscope. To collect T cells, grafts were mechanically dissociated by pipetting in FACS buffer. Single cell suspensions were then stained with Zombie NIR (BioLegend, Catalog No. 423105, 1/1000) for 30 min at 4° C. Samples were then washed with FACS buffer and incubated with the following Lineage antibodies for 30 min at 4°C: CD11b Biotin (BioLegend, Catalog No. 101204, 1/1000), CD11c Biotin (BioLegend, Catalog No. 117304, 1/1000), CD19 Biotin (BioLegend, Catalog No. 101504, 1/1000), DX5 Biotin (BioLegend, Catalog No. 108904, 1/1000), MHCII Biotin (BioLegend, Catalog No. 116404, 1/1000), GR1 Biotin (BioLegend, Catalog No. 108404, 1/1000), F4/80 Biotin (BioLegend, Catalog No. 123100, 1/1000), Ter119 Biotin (BioLegend, Catalog No. 116204, 1/1000) and NK-1.1 Biotin (BioLegend, Catalog No. 108704, 1/1000). After washes, samples were incubated with the following antibodies for 30 min at 4°C: CD45.1-PerCP-Cy5.5 (BioLegend, Catalog No. 110728, 1/500), CD45.2-BV650 (BioLegend, Catalog No. 109836, 1/200), CD4-BUV563 (Thermo Fisher Scientific, Catalog No. 365-0042-82, 1/1000), CD8-BUV615 (Thermo Fisher Scientific, Catalog No. 366-0081-82, 1/500), TCRαβ-PE-Dazzle594 (BioLegend, Catalog No. 109220, 1/500), TCRγδ-PE (BioLegend, Catalog No. 118108, 1/500), CD69-FITC (BioLegend, Catalog No. 104506, 1/500), CD24-APC (BioLegend, Catalog No. 101814, 1/1000), CD44-BV785 (BioLegend, Catalog No. 103059, 1/500), Ckit-BUV737 (Thermo Fisher Scientific, Catalog No. 367-1171-82, 1/200), CD71-PE-Cy7 (BioLegend, Catalog No. 113812, 1/200), Sca1-BUV395 (Thermo Fisher Scientific, Catalog No. 363-5981-82, 1/500), CD25-BV605 (BioLegend, Catalog No. 102036, 1/500), CD5-APC-eF780 (Thermo Fischer Scientific, Catalog No. 47-0015-82, 1/500) and Streptavidin-BV510 (Biolegend, Catalog No. 405234, 1/500). After final washes, samples were resuspended in FACS buffer and analyzed on an Aurora flow cytometer (Cytek Biosciences). Flow cytometry data were then analyzed using FlowJo (version 10.9.0).

### Sectioning, immunofluorescence staining and RNA scope on sections

Organoids, ORFTOCs, FTOCs and grafts were fixed in 4% paraformaldehyde in PBS for 30 min at room temperature (organoids) to overnight at 4 °C (ORFTOCs, FTOCs, grafts). Samples were then washed with PBS and either processed for cryosectioning or for paraffin embedding. For cryosectioning, samples were incubated in 30% (W/V) sucrose (Sigma-Aldrich, Catalog No. S1888) in PBS until the sample sank. Subsequently, samples were incubated for 12 h in a mixture of Cryomatrix (Epredia, Catalog No. 6769006) and 30% sucrose (Sigma-Aldrich, Catalog No. 84097) (mixing ratio 50/50), followed by a 12 h incubation in pure Cryomatrix. The samples were then embedded in a tissue mold, frozen on dry ice or in isopentane cooled by surrounding liquid nitrogen. 10 µm-thick sections were cut at -20°C using a CM3050S cryostat (Leica).

For paraffin embedding, organoid, ORFTOC and FTOC samples were embedded in HistoGel (Thermo Fisher Scientific, Catalog No. HG-4000-012) before being placed into histology cassettes. Cassettes were then processed with a Tissue-Tek VIP 6 AI Vacuum Infiltration Processor (Sakura) and embedded in paraffin. 4 µm paraffin sections were obtained with a Leica RM2265 microtome. Slides were processed through de-waxing and antigen retrieval in citrate buffer at pH 6.0 using a heat-induced epitope retrieval PT module (Thermo Fisher Scientific) before proceeding with immunostaining. Sections were then blocked and permeabilized for 30 min in 1% BSA (Thermo Fischer Scientific, Catalog No. 15260-037), 0.2% Triton X100 in PBS and blocked for 30 min in 10% goat or donkey serum in PBS at room temperature. Primary antibodies were incubated O/N at 4 °C in PBS, 1.5% donkey or goat serum. On the following day, slices were washed twice in 1% BSA, 0.2% Triton X-100 in PBS and incubated with secondary antibodies at room temperature for 45 min. Finally, slices were washed twice in 0.2% Triton X-100 in PBS and mounted with Fluoromount-G. The following primary and secondary antibodies were used: UEA1 (Vector Laboratories, Catalog No. B-1065, 1/500), Keratin 5 (BioLegend, Catalog No. 905501, 1/200), Keratin 8 (Abcam, Catalog. No. ab53280, 1/200), CD3ε (Thermo Fisher Scientific, Catalog No. MA5-14524, 1/200), MHCII-Biotin (BioLegend, Catalog No. 107603, 1/200) EpCAM-PE (BioLegend, Catalog No. 118206, 1/200), Aire (Thermo Fischer Scientific, Catalog No. 14-5934-82, 1/50), Streptavidin Alexa 488 (Thermo Fisher Scientific, Catalog No. S-11223, 1/500), Goat anti-Rat Alexa 568 (Thermo Fisher Scientific, Catalog No. A-11077, 1/500), and Donkey anti-Rabbit Alexa 647 (Thermo Fisher Scientific, Catalog No. A-31573, 1/500). Nuclei were again stained with Dapi (Tocris, Catalog No. 4748, 1 ug/ml). RNAscope Multiplex Fluorescent V2 assay (Bio-Techne, catalog no. 323110) was performed according to the manufacturer’s protocol. Paraffin sections were hybridized with the probes Mm-Foxn1 (Bio-Techne, catalog no. 482021). Mm-3Plex probes (Bio-Techne, catalog no. 320881) and 3Plex Dapb probes (Bio-Techne, catalog no. 320871) were used as positive and negative controls, respectively. Probes were incubated at 40°C for 2 hours, and the different channels were revealed with TSA Opal570 (Akoya Biosciences, catalog no. FP1488001KT). Tissues were counterstained with Dapi and mounted with ProLong Gold Antifade Mountant (Thermo Fisher Scientific, P36930). Hematoxylin and eosin staining was performed using a Ventana Discovery Ultra automated slide preparation system (Roche).

### Microscopy and image analysis

Live brightfield imaging was performed using a Nikon Eclipse Ti2 inverted microscope with 4×/0.13 NA, 10×/0.30 NA, and 40×/0.3 NA air objectives and a DS-Qi2 camera (Nikon Corporation). Time lapse was imaged with a Nikon Eclipse Ti inverted microscope system equipped with a 20×/0.45 NA air objective and a DS-Qi2 camera (Nikon Corporation). Both microscopes were controlled using the NIS-Elements AR software (Nikon Corporation). Extended depth of field (EDF) of brightfield images was calculated using a built-in NIS-Elements function. Fluorescent confocal imaging of fixed whole-mount and sections was done on a Leica SP8 microscope system, equipped with a 20×/0.75 NA air and a 40x/1.25 glycerol objectives, 405 nm, 488 nm, 552nm and 638 nm solid state lasers, DAPI, FITC, RHOD and Y5 filter cubes, a DFC 7000 GT (Black/White) camera and a CCD grayscale chip. Sections were also imaged on a Leica DM5500 upright microscope equipped with a 20x/0.7 NA air and a 40x/1 NA oil objectives, a DFC 3000 (Black/White) or a DMC 2900 (Color) cameras and a CCD grayscale or a CMOS color chip, respectively. Both Leica microscopes were controlled by the Leica LAS-X software (Leica microsystems). For image processing, only standard contrast- and intensity-level adjustments were performed, using Fiji/ImageJ (NIH) (version 2.1.0/1.53c).

### Statistics

The number of replicates (*n*), the number of independent experiments or animals, the type of statistical tests performed, and the statistical significance are indicated for each graph in the figure legend. Statistical significance was analyzed using one- or two-way ANOVA, Brown-Forsythe ANOVA in case of heteroscedasticity or Mood’s median test in the absence of normal distribution. For multiple comparisons, one-way ANOVA were followed by Tukey’s test, Brown-Forsythe ANOVA by Dunnet’s T3 test, and Mood’s test results adjusted for false-discovery rate. Data normality and equality of variances were previously tested with Shapiro-Wilk and Brown-Forsythe test, respectively. Grubbs test was used to determine the presence of outliers across scRNAseq subpopulations. In all cases, values were considered significant when P ≤ 0.05. Graphs show individual datapoints with mean ± standard deviation (SD). Tests were performed using Prism (GraphPad, version 9.4.0), except Grubbs test which was performed using GraphPad website (https://www.graphpad.com/quickcalcs/grubbs1/) and Mood’s test which was performed using the package rcompanion (Mangiafico, 2016) in R (version 4.1.2). Graphs were made using Prism.

## Acknowledgements

We thank all present and past LSCB members and in particular Antonius Chrisnandy, Bilge Sen Elci and Moritz Hofer for discussions and sharing materials. We thank Julia Prébandier for administrative assistance and Katrin Hafen for technical expertise. We acknowledge support and work from the Flow Cytometry, Centre de Phénogénomique, Gene Expression, Histology and BioImaging and Optics EPFL core facilities. We thank all collaborators from the Syn-Thy project, in particular Graham Anderson, Viktorja Major, Joanna Sweetman and Tim Henderson and for their feedback.

## Author contributions

TH conceived the study, designed and carried experiments, analyzed results, prepared artwork, and wrote the manuscript. LFLM helped with experimental design, experimental and analysis work and provided feedback on the manuscript. TB performed grafting experiment and analysis and provided feedback. LT and JJL helped with experiments. PR taught methods, provided feedback and shared reagents. CCB and GH provided feedback on the work and edited the manuscript. MPL conceived the work, designed experiments, and carried out the final editing of the manuscript.

## Data availability

Sequencing data reported in this paper have been deposited in the Gene Expression Omnibus (GEO) public repository under the accession number GSE240698. The SubSeries GSE240696 and GSE240697 corresponds to bulk and single-cell RNA-seq, respectively. Code used for analysis is available upon request and will be published on Github.

## Supplementary information

**Fig. S1.**
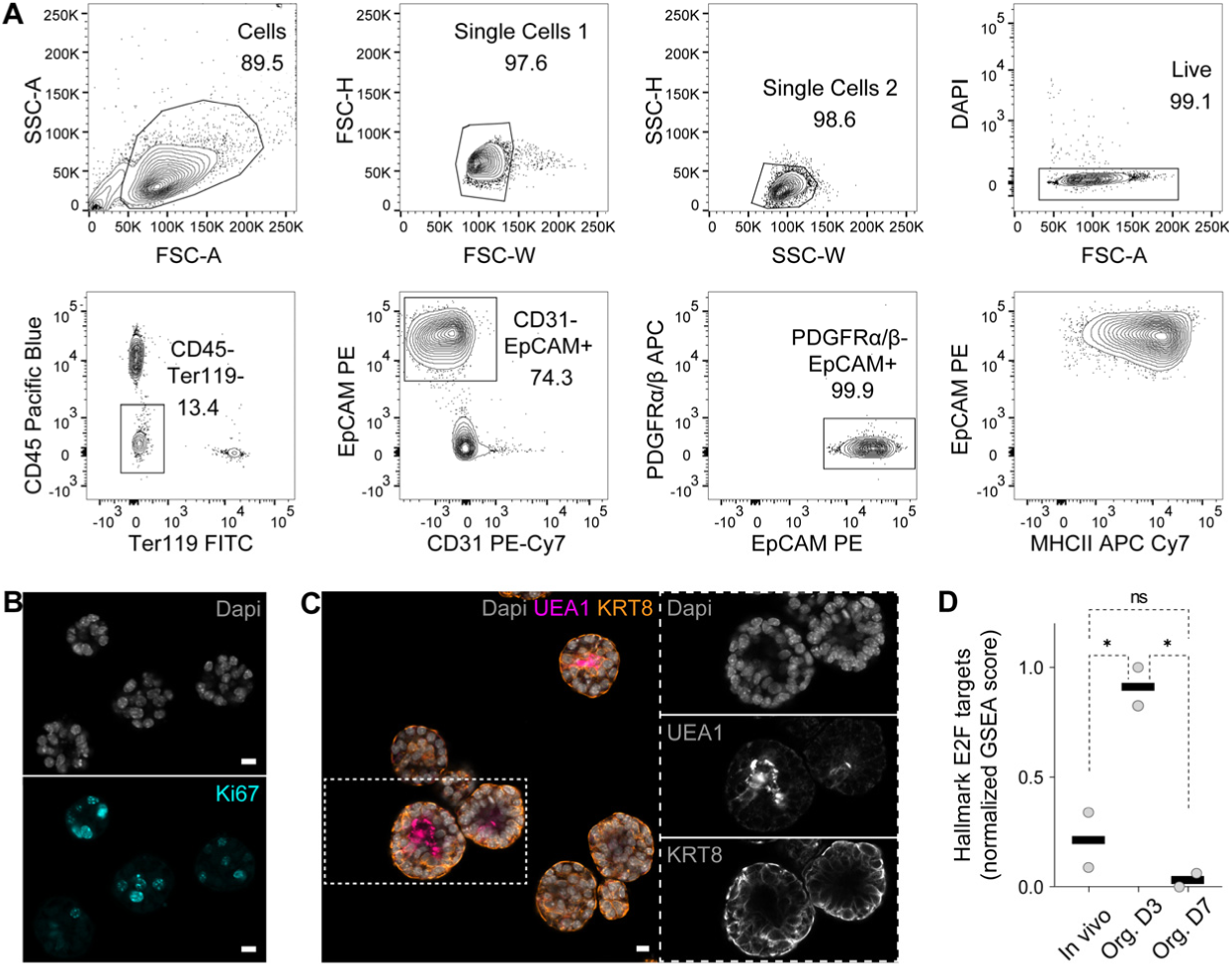
Establishment of thymic epithelial organoids. (**A**) Sorting strategy to isolate TECs (EpCAM+). Exclusion of CD45+, Ter119+, CD31+ and PDGFRα+ PDGFRβ+ cells. TECs are mostly MHCII+. (**B**) Immunofluorescence images of D7 organoids showing nuclei (Dapi, top [grey]) and proliferating cells (Ki67, bottom [cyan]). Scale bars, 10μm. (**C**) Immunofluorescence image of D7 organoids highlighting different cell populations, here with medullary cells (UEA1-reactivity [bright pink and middle] and cortical cells (KRT8 [amber and bottom]). Dapi counterstains nuclei (grey and top). Scale bar, 10μm. (**D**) Single-sample gene set enrichment analysis (GSEA) demonstrating a proliferation peak (E2F hallmark) for D3 organoids. ns: P > 0.05, *: P = 0.0241 (In vivo vs Org. D3) and P = 0.0126 (Org. D3 vs Org. D7) (one-way ANOVA with Tukey’s multiple comparisons test; n = 2 mice per condition). Graph represents individual datapoints with mean.

**Fig. S2.**
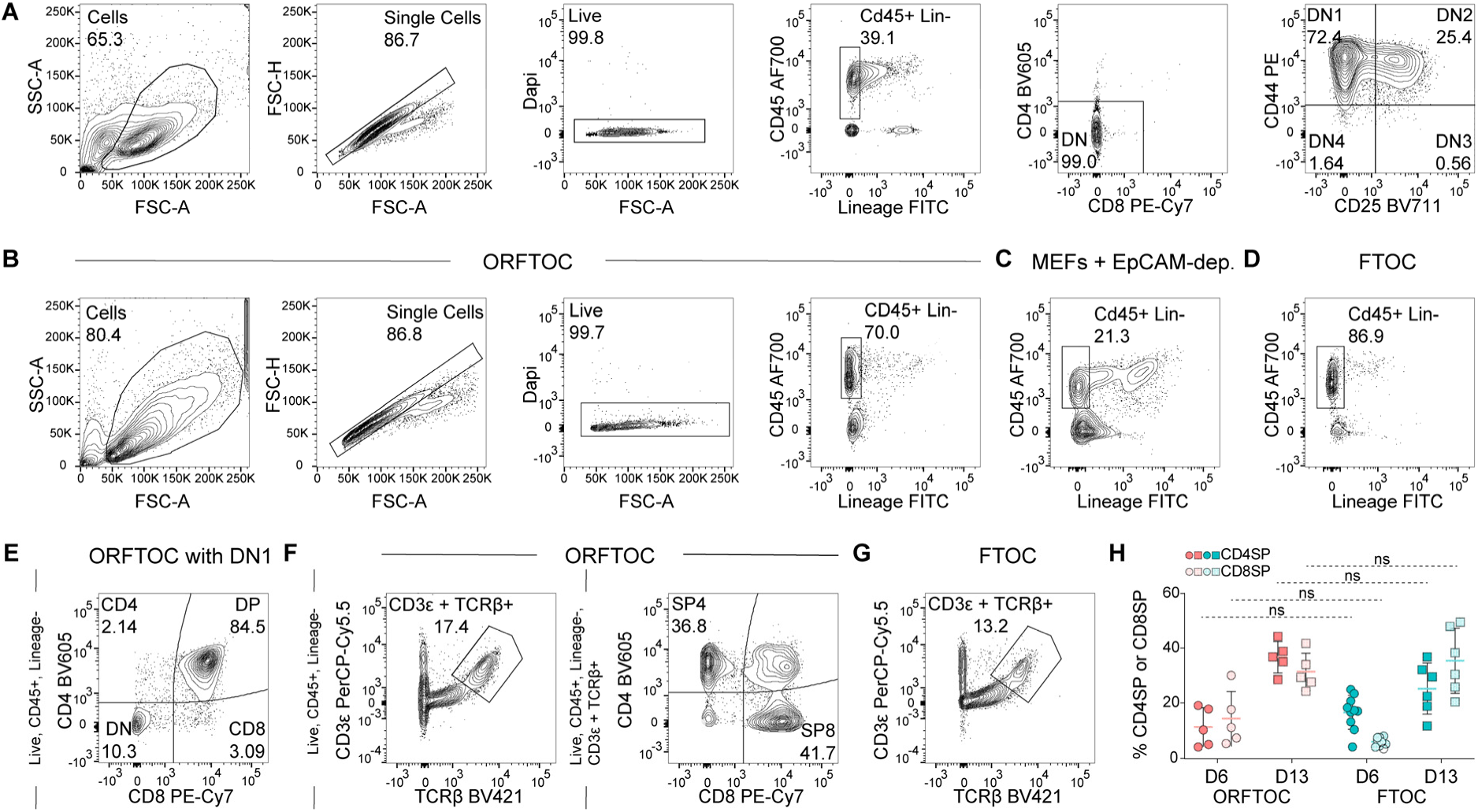
Generation and analysis of ORFTOCs and control conditions. (**A**) Flow cytometry plots showing the developmental stage (DN) of thymocytes in E13.5 thymi. (**B**) Flow cytometry plots demonstrating the gating strategy used to analyze thymocyte development in ORFTOCs and control conditions. The last plot highlights the CD45+ Lin-population in ORFTOCs at D6. (**C-D**) Flow cytometry plots representing the CD45+ Lineage-population in control reaggregates with MEFs and the EpCAM-depleted fraction of cells from E13.5 thymi (C) and in FTOC controls (D) at D6. (**E**) Flow cytometry plot showing T cell development in D13 ORFTOCs made with adult DN1 as T cell input population. Gating strategy is indicated on the left. (**F**) Flow cytometry plots representing the CD3ε+ TCRβ+ population in D13 ORFTOCs, and its division into CD4SP and CD8SP T cell lineages, with gating strategies indicated on the left of each plot. (**G**) Flow cytometry plot showing the CD3ε+ TCRβ+ population in D13 FTOC controls. Gating strategy is as in F (left). (**H**) Percentage of CD4SP and CD8SP within the CD3ε+ TCRβ+ population, at D6 and D13, for both ORFTOCs and FTOC controls. ns: P > 0.05 (Brown-Forsythe ANOVA test for both CD4SP and CD8SP with Dunnet’s T3 multiple comparisons test; n = 5 for all ORFTOC populations and time, n = 10 for FTOC populations at D6 and n = 6 for FTOC populations at D13, from 5 independent experiments). Graph represents individual datapoints with mean + SD.

**Fig. S3.**
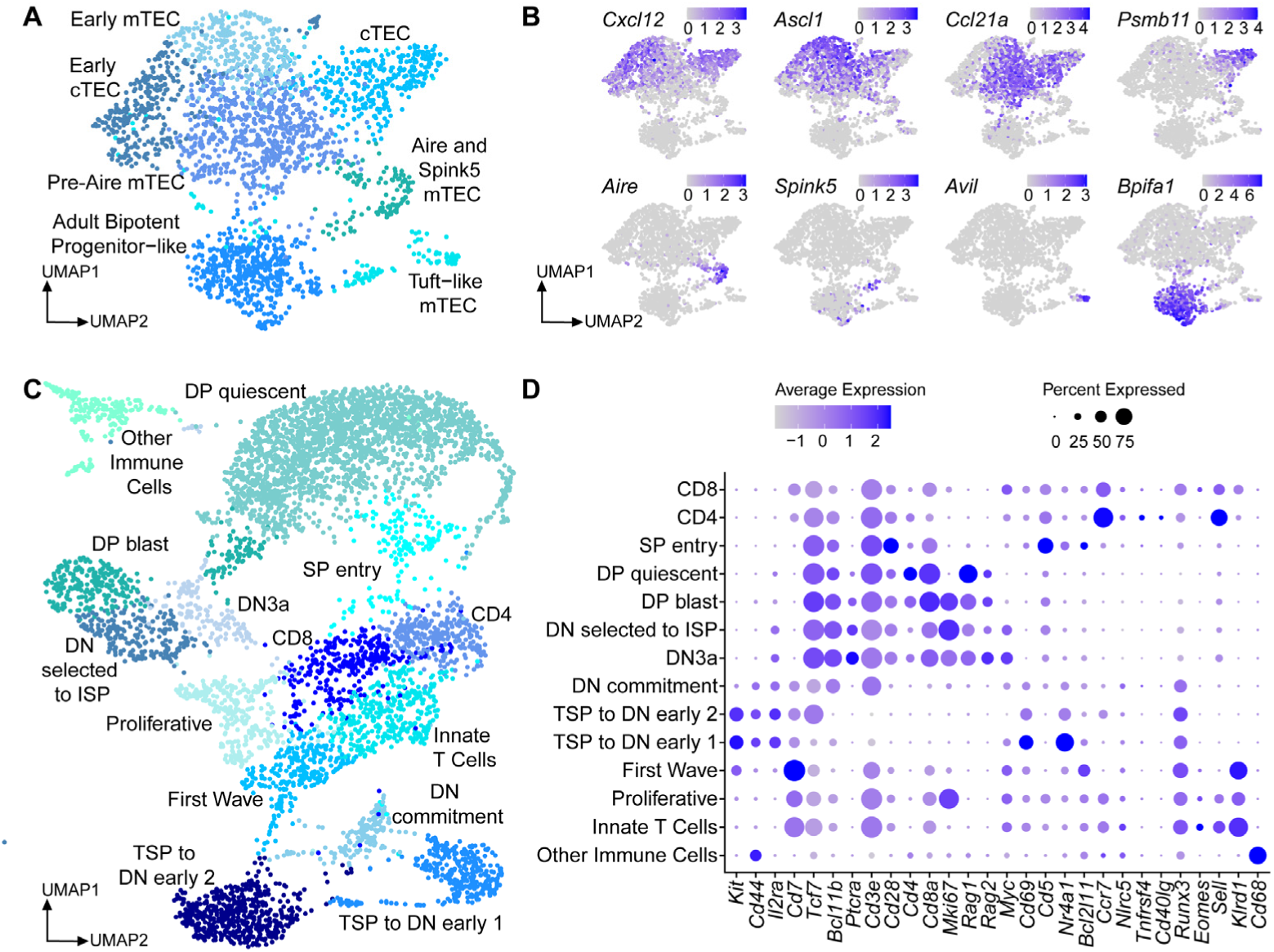
Single-cell transcriptomic characterization of ORFTOCs and FTOCs. (**A**) UMAP showing the different epithelial clusters. (**B**) UMAPs highlighting characteristic marker expression for each of the epithelial clusters. (**C**) UMAP showing the different immune clusters. (**D**) Dot plot summarizing the expression of characteristic markers of T cell development for the different clusters.

**Fig. S4.**
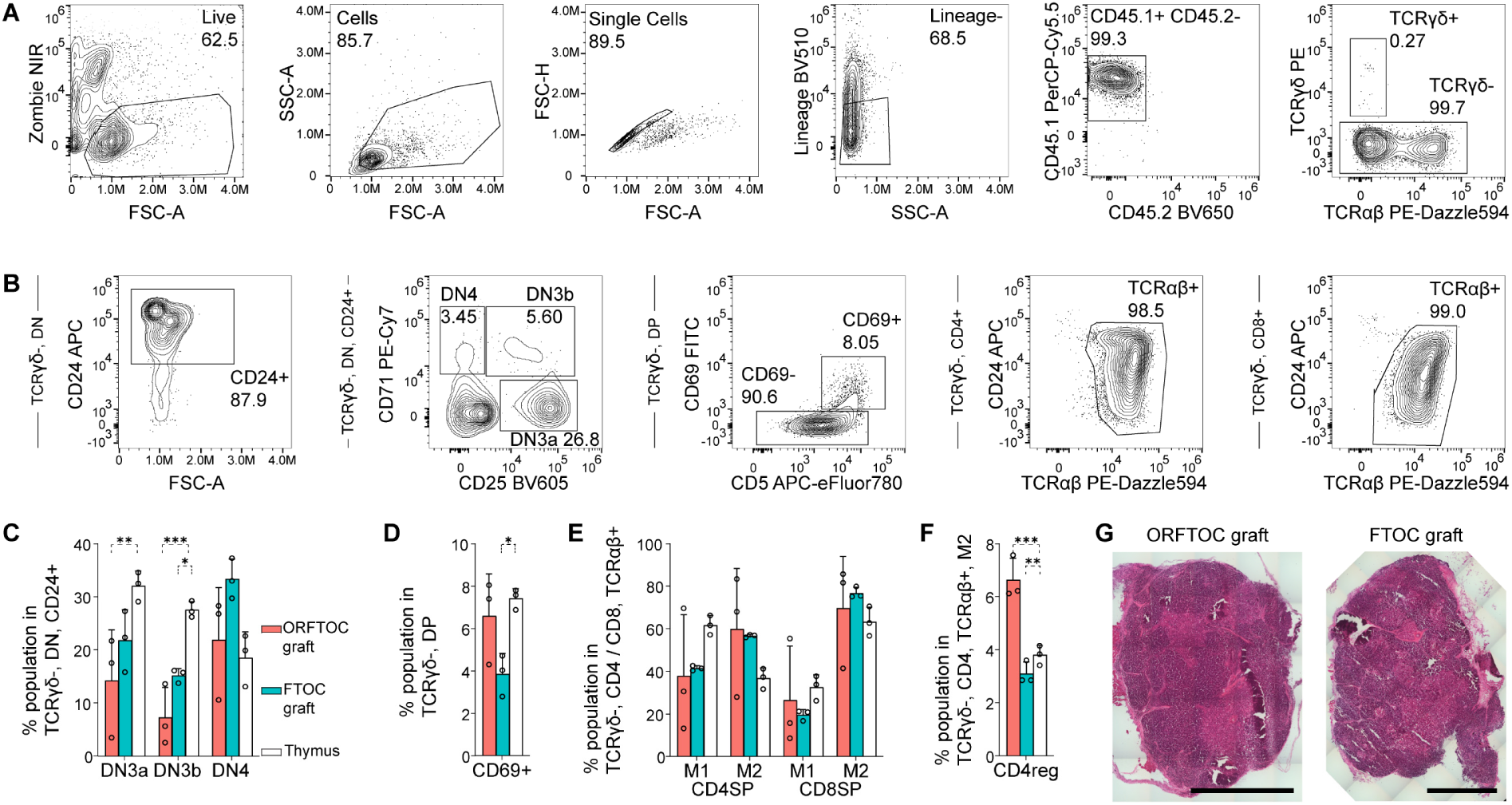
Analysis of ORFTOC grafts and comparison to controls. (**A - F**) Flow cytometry plots showing the gating strategy to analyze grafts. ORFTOCs (CD45.2) were grafted under the kidney capsule of CD45.1 hosts. Live, single, Lineage negative (CD11b, CD11c, Gr1, Ter119, DX5, NK-1.1, MHCII, F4/80) cells were gated for CD45.1 positivity (A) and TCRγδ negative T cells were further analyzed (B). CD25 and CD71 expression on CD24+ DN cells were used to enumerate DN3a, DN3b and DN4 subsets, quantified in (C) for the different conditions. β-selection occurs at the DN3a to DN3b transition. CD69 expression identifies cells undergoing positive selection and is quantified in (D) for the different conditions. Gating on the TCRαβ+ population, mature (M1 and M2) CD4SP and CD8SP T cells are quantified in E for the different conditions. Within the M2 population, CD4 regulatory T cells (CD4reg) are quantified in F for the different conditions. For all bar graphs, only significant differences are indicated with stars. * P = 0.0381 (DN3a ORFTOC vs thymus), *** P = 0.0009 (DN3b ORFTOC vs thymus), * P = 0.0116 (DN3b FTOC vs thymus), * P = 0.0373 (Cd69 FTOC vs thymus), ns: P > 0.05 (one-way ANOVA for each subpopulation between conditions, n = 3 grafts/mice for each condition). Bar graphs represent mean and SD, with individual datapoints displayed as circles. (**G**) Hematoxylin and eosin (H&E) staining of ORFTOC and FTOC grafts.

**Movie 1. Thymic epithelial organoid establishment.** One week time-lapse showing the development of thymic epithelial organoids starting from sorted single thymic epithelial cells seeded in Matrigel and cultured in defined conditions.

